# NBS1 binds directly to TOPBP1 via disparate interactions between the NBS1 BRCT1 domain and the TOPBP1 BRCT1 and BRCT2 domains

**DOI:** 10.1101/2022.09.26.509489

**Authors:** Oanh Huynh, Kenna Ruis, Katrina Montales, W. Matthew Michael

## Abstract

The TOPBP1 and NBS1 proteins are key components of DNA repair and DNA-based signaling systems. TOPBP1 is a multi-BRCT domain containing protein that plays important roles in checkpoint signaling, DNA replication, and DNA repair. Likewise, NBS1, which is a component of the MRE11-RAD50-NBS1 (MRN) complex, functions in both checkpoint signaling and DNA repair. NBS1 also contains BRCT domains, and previous works have shown that TOPBP1 and NBS1 interact with one another. In this work we examine the interaction between TOPBP1 and NBS1 in detail. We report that NBS1 uses its BRCT1 domain to interact with TOPBP1’s BRCT1 domain and, separately, with TOPBP1’s BRCT2 domain. Thus, NBS1 can make two distinct contacts with TOPBP1. We report that recombinant TOPBP1 and NBS1 proteins bind one another in a purified system, showing that the interaction is direct and does not require post-translational modifications. Surprisingly, we also report that intact BRCT domains are not required for these interactions, as truncated versions of the domains are sufficient to confer binding. For TOPBP1, we find that small 24-29 amino acid sequences within BRCT1 or BRCT2 allow binding to NBS1, in a transferrable manner. These data expand our knowledge of how the crucial DNA damage response proteins TOPBP1 and NBS1 interact with one another and set the stage for functional analysis of the two disparate binding sites for NBS1 on TOPBP1.

## 1. Introduction

The BRCA1 C-terminal (BRCT) domain is an interaction module that allows the host protein to bind to various biological substrates such as other proteins, nucleic acids, and poly (ADP-ribose) chains (reviewed by Gerloff et al, 2012; Leung and Glover, 2011; Reinhardt and Yaffe, 2013; Wan et al, 2016). BRCT proteins are major players in the DNA damage response (DDR), and it is therefore crucial that we understand how they function and that we fully describe all the different modes by which they interact with their binding partners. Regarding how BRCT domains bind to other proteins, there are two general categories: interactions that require the BRCT binding substrate to be phosphorylated, and those that do not. For phospho-driven interactions, some BRCT domains contain a so-called phosphate-binding ocket (PBP), which features a triad of conserved amino acid residues that interact with the phosphate group on the substrate protein (reviewed by Leung and Glover, 2011; Reinhardt and Yaffe, 2013; Wan et al, 2016; Day et al, 2021). BRCT domains that bind phospho-epitopes are typically found in pairs, where one domain within the pair contains the PBP. While the PBP directly interacts with phosphate, both BRCT domains within the pair are required for binding as the PBP forms via intramolecular interactions between the two domains (reviewed by Leung and Glover, 2011; Reinhardt and Yaffee, 2013; Day et al, 2021). Intramolecular interactions between BRCT domains also allow binding of the host protein to substrate in a phospho-independent manner, and both the phospho-dependent and independent modes of binding have been observed within the same protein. The yeast Rtt107 protein contains 6 BRCT domains, with the C-terminal two BRCTs supplying phospho-dependent binding to histone H2A, and the four N-terminal domains forming a compact, higher-order assembly that binds peptide ligands from a number of proteins (Wan et al, 2019). The net result is that Rtt107 binds to chromatin via the phospho-dependent mechanism, and then recruits other proteins to chromatin via the phospho-independent mechanism.

BRCT domains also make intermolecular contacts to allow assembly of both homo- and heterotypic multimers. A well-studied example of BRCT::BRCT intermolecular interaction is found with the XRCC1 and DNA Ligase 3 (LIG3) proteins (Taylor et al, 1998; Dulic et al, 2001; Beernink et al, 2005; Cuneo et al, 2011; London, 2015; London et al, 2020; Hammel et al, 2021). Both proteins can homodimerize via BRCT::BRCT interaction, and the two can also heterodimerize via their BRCT domains. Recent work from our group was focused on the Topoisomerase II Binding Protein 1 (TOPBP1) protein, which contains nine BRCT domains, and we found that TOPBP1 forms oligomers and that the protein is capable of a series of BRCT::BRCT interactions (Kim et al, 2020). More specifically, we found that BRCT1&2 could bind to BRCT domains 1,2, and 5, and that BRCT4&5 could bind BRCTs 1 and 2 but not 5. In this work we examine how TOPBP1 uses its BRCT domains to bind a heterologous protein, by studying its interaction with the NBS1 protein.

NBS1 is part of a trimeric complex, termed MRN for MRE11-RAD50-NBS1, that senses DNA double-strand breaks (DSBs) and then plays multiple roles in ensuring efficient repair of the breaks (reviewed by McCarthy-Leo et al, 2022; Reginato and Cejka, 2020; Syed and Tainer, 2018). These roles include holding the two broken ends together, initiating DNA end resection, and activation of the DNA damage checkpoint kinases ATM and ATR. MRN activates ATM directly (reviewed by Paull, 2015), and ATR indirectly through its ability to recruit TOPBP1 to the DSB (Montales et al, 2022). Recent work from our laboratory has shown that loss of MRN attenuates ATR signaling at DSBs, and TOPBP1’s occupancy at the DSB is also reduced (Montales et al, 2022). TOPBP1 has a well-studied function in directly interacting with ATR (and its binding partner ATRIP) at sites of damage to stimulate ATR kinase activity (reviewed by Wardlaw et al, 2015), and thus our recent work has revealed a pathway for ATR signaling that runs from the DNA end through MRN to TOPBP1 and then ATR (Montales et al, 2022). One open and important question for this pathway is: what is the molecular mechanism by which MRN recruits TOPBP1 to DSBs? In this study we focused on the NBS1 subunit as previous works have shown that TOPBP1 and NBS1 bind one another, and that this interaction is conserved from frogs to man (Morishima et al, 2007; Yoo et al, 2009). Interestingly, in both frogs and humans, TOPBP1 and NBS1 constitutively bind to one another, and the interaction is then enhanced during the DSB response (Morishima et al, 2007; Yoo et al, 2009). Here, we study the constitutive interaction, and we find that NBS1 uses its BRCT1 domain to make direct contacts with TOPBP1’s BRCT1 domain and, separately, the BRCT2 domain. Our data provide a molecular explanation for how MRN recruits TOPBP1 to DSBs, and they also expand our understanding of how BRCT::BRCT interactions drive the cellular response to DNA damage.

## 2. Material and Methods

### 2.1 Materials

#### 2.1.1 Plasmids

The expression vector to produce GST-RAD9 Tail has previously been described (Duursma et al., 2013). All other plasmids used in this study are described below, in table format, and all are *Xenopus* proteins. Specific details about plasmid construction are available upon request.

| Plasmid | Name | Vector | AA coordinates |
| --- | --- | --- | --- |
| 5Xmyc-BRCT0-2 | pHG128 | pCS2+MT | 1-333 |
| 5Xmyc-BRCT0-2 KK155,249EE | pAK42 | pCS2+MT | 1-333 |
| 5Xmyc-BRCT4&5 | pJA19 | pCS2+MT | 480-758 |
| 5Xmyc-BRCT0 | pHG126 | pCS2+MT | 1-99 |
| 5Xmyc-BRCT1 | pHG129 | pCS2+MT | 100-191 |
| 5Xmyc-BRCT2 | pJA23 | pCS2+MT | 191-333 |
| 5Xmyc-NBS1 | pKM107 | pCS2+MT | 1-763 |
| 5Xmyc-NBS1 1-400 | pKM105 | pCS2+MT | 1-400 |
| 5Xmyc-NBS1 1-200 | pKR34 | pCS2+MT | 1-200 |
| 5Xmyc-NBS1 1-100 | pOH4 | pCS2+MT | 1-100 |
| 5Xmyc-NBS1 100-200 | pKR36 | pCS2+MT | 100-200 |
| 5Xmyc-NBS1 200-400 | pKR37 | pCS2+MT | 200-400 |
| 5Xmyc-NBS1 393-763 | pKM106 | pCS2+MT | 393-763 |
| 5Xmyc-NBS1 1-400 K158A | pOH34 | pCS2+MT | 1-400 |
| 5Xmyc-NBS1 1-175 | pOH35 | pCS2+MT | 1-175 |
| 5Xmyc-NBS1 1-150 | pOH36 | pCS2+MT | 1-150 |
| 5Xmyc-NBS1 128-400 | pOH39 | pCS2+MT | 128-400 |
| 5Xmyc-NBS1 128-400 | pOH40 | pCS2+MT | 150-400 |
| MBP-TOPBP1 1-333 | pAK53 | pUC57-MBP | 1-333 |
| MBP-TOPBP1 91-275 | pAK56 | pUC57-MBP | 91-275 |
| MBP-TOPBP1 91-196 | pKR23 | pUC57-MBP | 91-196 |
| MBP-TOPBP1 91-179 | pKR24 | pUC57-MBP | 91-179 |
| MBP-TOPBP1 91-156 | pKR25 | pUC57-MBP | 91-156 |
| MBP-TOPBP1 91-139 | pKR26 | pUC57-MBP | 91-139 |
| MBP-TOPBP1 128-179 | pKR29 | pUC57-MBP | 128-179 |
| MBP-TOPBP1 138-179 | pKR30 | pUC57-MBP | 138-179 |
| MBP-TOPBP1 128-156 | pKR38 | pUC57-MBP | 128-156 |
| MBP | pOH10 | pUC57-MBP | - |
| MBP-TOPBP1 221-273 | pOH6 | pUC57-MBP | 221-273 |
| MBP-TOPBP1 231-273 | pOH7 | pUC57-MBP | 231-273 |
| MBP-TOPBP1 221-250 | pOH8 | pUC57-MBP | 221-250 |
| MBP-TOPBP1 197-231 | pOH9 | pUC57-MBP | 197-231 |
| MBP-TOPBP1 242-273 | pOH11 | pUC57-MBP | 421-273 |
| MBP-TOPBP1 250-273 | pOH12 | pUC57-MBP | 250-273 |

#### 2.1.2 Recombinant proteins

The recombinant proteins used in this study were GST, GST-NBS1 1-400, GST-NBS1 1-400 K158E, GST-NBS1 393-763, GST-RAD9Tail, GST-BRCT4&5, MBP, MBP-BRCT1&2, and 6His-T7-BRCT0-2. Expression vectors for GST-RAD9Tail (Duursma et al., 2013), GST-BRCT4&5, MBP-BRCT1&2, and 6His-T7-BRCT0-2 (Kim et al, 2020) have been described. Expression vectors for GST-NBS1 1-400, GST-NBS1 1-400 K158E, and GST-NBS1 393-763 were made using pGEX4T-3 as the backbone. All proteins were expressed in *E. coli* BL21(DE3) cells, at 37°C for 4 hours, and all proteins were purified from the soluble fraction according to standard procedures (detailed below).

#### 2.1.3 Antibodies

We used the following commercially sourced antibodies in this work: Myc (Millipore Sigma #M4439), MBP (New England Biolabs #E8032S), GST (Millipore Sigma #05-782), and T7 (Millipore Sigma #69522). We also used our own antibody against *Xenopus* TOPBP1, HU142, which has been described (Van Hatten et al., 2002).

### 2.2 Methods

#### 2.2.1 Xenopus egg extracts (XEE)

Extracts were prepared as previously described (Van Hatten et al., 2002, Yan et al., 2006).

#### 2.2.2 IVTT production of proteins and determination of protein concentrations

In-vitro transcription and translation (IVTT) reactions were performed using the SP6 TnT® Quick Coupled Transcription/Translation System (Promega #L2080) according to the manufacturer’s instructions. For Bug-Bunny binding assays we used 20µl of the IVTT reaction/sample.

#### 2.2.3 Recombinant Protein Expression and Purification

Plasmids were transformed in BL21 (DE3) Competent *E. coli* cells and grown in LB media (GST fusion proteins) or LB media with 0.2% glucose (MBP fusion proteins) with ampicillin. At an OD600 of 0.6-0.8, cells were induced with 0.3 mM IPTG for 4 hours at 37 °C. Cells were then harvested by centrifugation (4000 rpm for 20 min) and washed with phosphate-buffered saline (PBS). Cells were lysed by sonication in GST column buffer (40% PBS, 60% TEN buffer: 10 mM Tris-Cl, pH 8.0, 1 mM EDTA, pH 8.0, 100 mM NaCl) supplemented with complete protease inhibitor cocktail tablets (Roche #04693159001) and 10 mg lysozyme for GST fusion proteins or in maltose column buffer (20 mM Tris-HCl, pH 7.4, 0.3 M NaCl, and 1 mM EDTA) with 1mM DTT and complete protease inhibitor cocktail tablets for MBP fusion proteins. Lysates were cleared by centrifugation (15,000 rcf for 20 min) and incubated with glutathione sepharose (BioVision #6566) for GST fusion proteins or amylose resin (New England Biolabs #E8021S) for MBP fusion proteins for 1 hour at 4°C. Beads were poured over a column and washed with column buffer. GST fusion proteins were eluted with 20 mM glutathione in 75 mM HEPES, pH 7.5, 150 mM NaCl, 5 mM DTT, and 0.1% Triton-X. MBP fusion proteins were eluted with 10 mM maltose in maltose column buffer. Proteins were dialyzed overnight into PBS.

#### 2.2.4 Protein binding assays

Three kinds of protein binding assays were utilized. The first involved the addition of recombinant proteins to XEE (Figures 1B-D). For these assays 25 µl of XEE was supplemented with 5 µg of GST-tagged recombinant protein. DSBs, in the form of EcoRI-digested lambda DNA, were also optionally added (500 ng). After a 30-minute incubation the samples were diluted into 400 µl of Binding Buffer (PBS + 0.1% NP-40) and incubated with glutathione sepharose (BioVision, #6566). The beads were washed four times with 500 µl of Binding Buffer and then eluted with 2X Sample Buffer (Millipore Sigma #S3401-10VL). The total and bound fractions were then analyzed via Western blotting using standard conditions. The second involved incubation of IVTT proteins with recombinant GST or MBP fusion proteins purified from *E. coli*, in what we refer to as the Bugs-Bunny system. For these assays 20 µl of IVTT proteins were mixed with 5 µg of GST or MBP fusion protein and incubated for 60 minutes at 30°C. The mixtures were diluted into 400 µl of Binding Buffer and then incubated with the appropriate resin (amylose for MBP, New England Biolabs # E8021S; glutathione sepharose for GST, BioVision #6566) for one hour at 4°C. The beads were washed four times with 500 µl of Binding Buffer and then eluted with 2X Sample Buffer. The input and bound fractions were then analyzed via Western blotting using standard conditions. The third type of binding assay (done once, in Figure 4F) was a variant of the pull-down where MBP recombinant proteins, purified from *E. coli*, were mixed at 10 µg each with 100 ng of 6His-T7-BRCT0-2, also purified from *E. coli*. The samples were then treated as described above for the Bugs-Bunny system. ImageJ software was used to quantify the signal intensity of bound and input samples, and the percent of bound to input was presented in Figure 2C.

**Figure 1.**
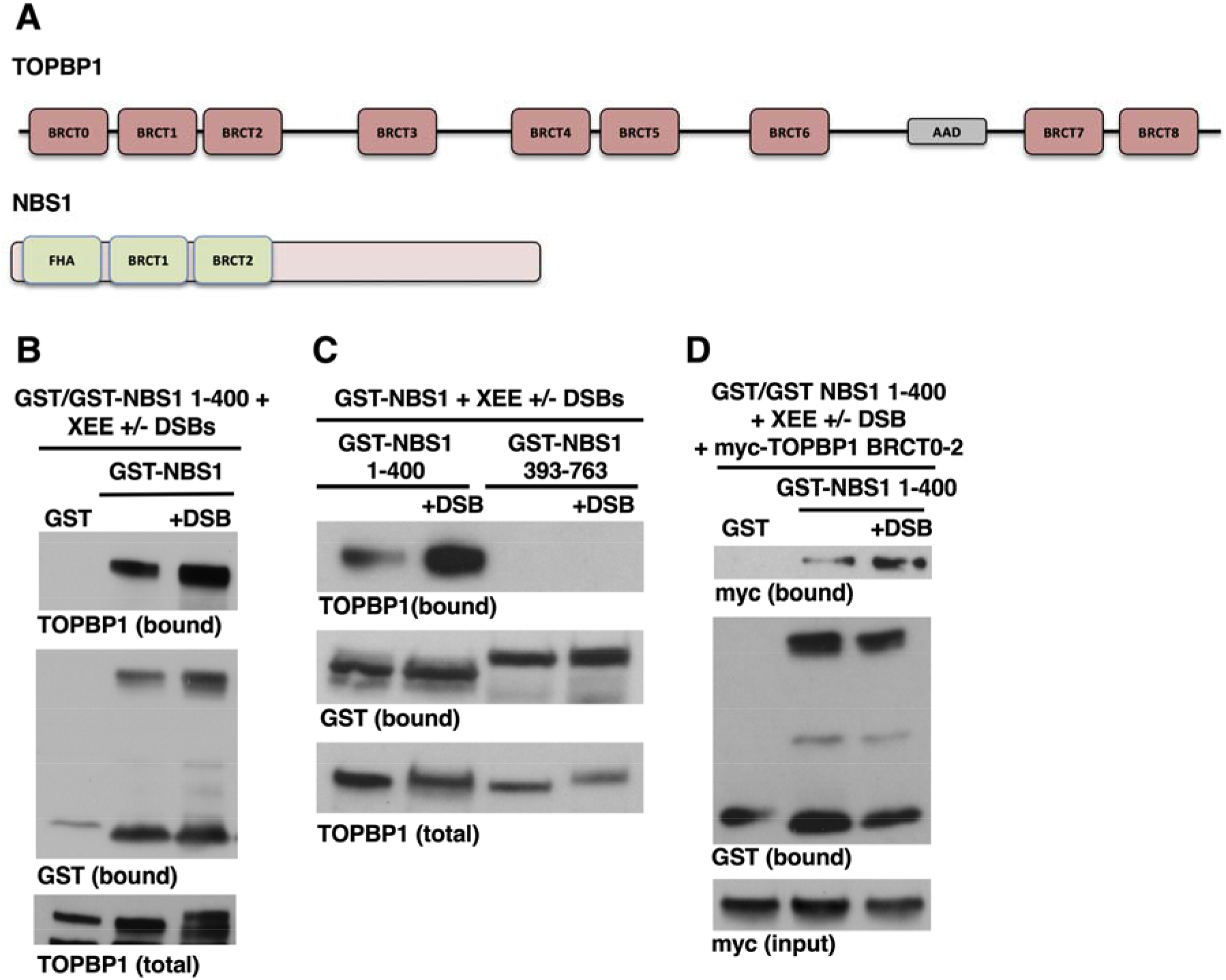
NBS1 and TOPBP1 interact in frog egg extracts. **A**. Schematic representation of the proteins of interest. TOPBP1 contains 9 BRCT domains while NBS1 contains two BRCT domains, as well as an N-terminal FHA domain. **B**. A GST pull-down assay was performed, as described in the Methods, with the indicated GST proteins. This was done in the presence of egg extract, with optionally added DSBs, and the presence of endogenous TOPBP1 in the bound fraction was detected by Western blot. The experiment shown is representative of two independently performed biological replicates. **C**. Same as (B) except GST-NBS1 393-763 was included. The experiment shown is representative of two independently performed biological replicates. **D**. Myc-tagged TOPBP1 BRCT0-2, produced by IVTT, was added to XEE, and DSBs were optionally included, as indicated. The indicated GST proteins were also included, and then the samples were then processed as in (B). The experiment shown is representative of two independently performed biological replicates.

**Figure 2.**
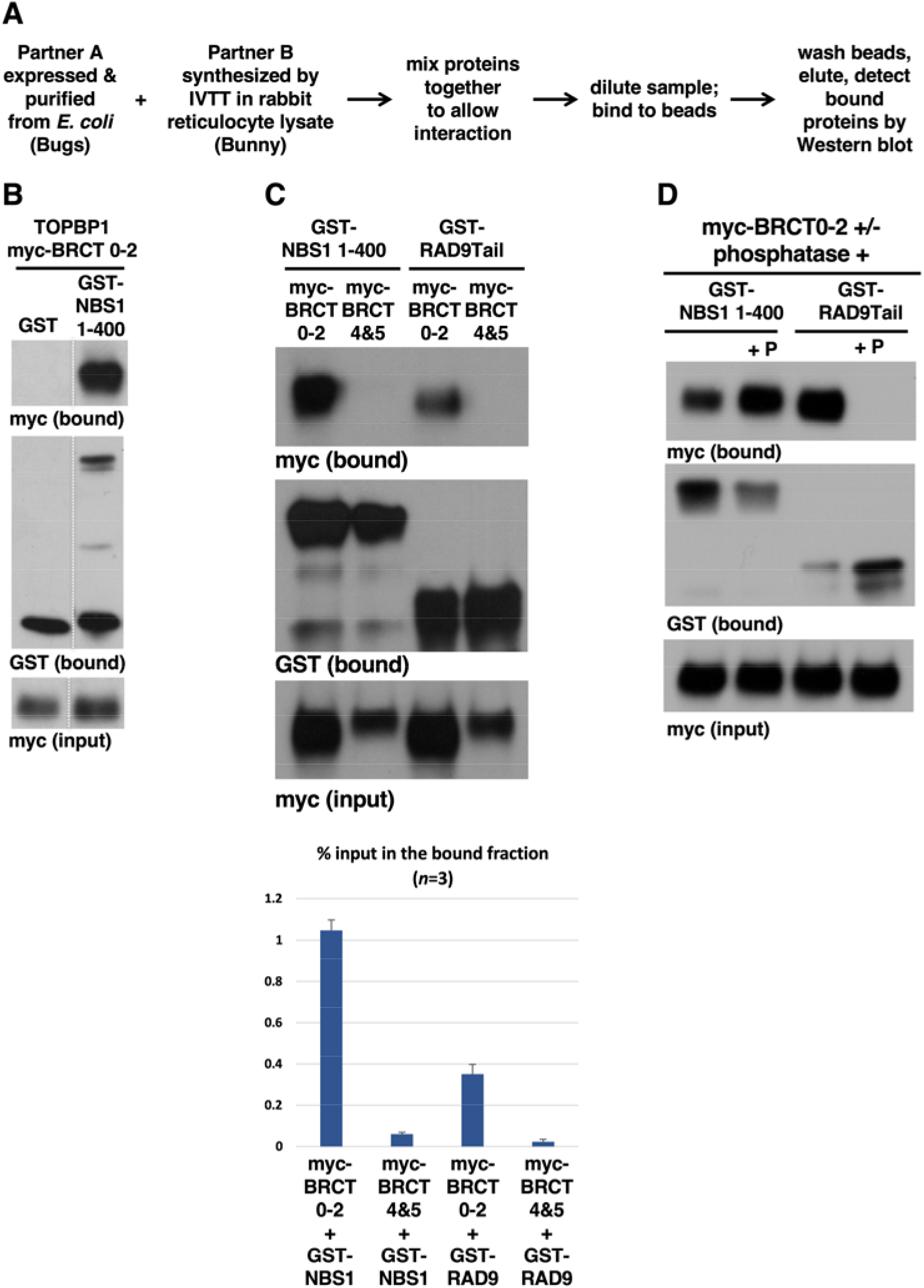
The Bugs-Bunny system recapitulates a phosphate-independent interaction between NBS1 and TOPBP1. **A**. Overview of the Bugs-Bunny protein binding assay system. **B**. The indicated GST proteins, expressed and purified from *E. coli*, were incubated in IVTT reactions expressing myc-tagged TOPBP1 BRCT0-2. The samples were then processed for the GST pull-down assay as described in the Methods. Samples of the total binding reaction, taken prior to dilution, were also probed for myc, showing that the myc-tagged proteins were present in equivalent amounts across all the samples (lane “input”). The experiment shown is representative of two independently performed biological replicates. We note that the image shown is derived from a larger gel, and the irrelevant samples were removed. The white dashed line depicts where the gel was spliced together. The full gel with all original samples is shown in Figure S1A. **C**. A GST pull-down assay was performed, as in part A, with the indicated GST- and myc-tagged proteins. The experiment shown is representative of three independently performed biological replicates. Quantification of all three experiments is shown in the graph. Please see Methods for how these experiments were quantified. **D**. A GST pull-down assay was performed, as in part A, with the indicated GST proteins and myc-tagged TOPBP1 BRCT0-2. In this experiment lambda phosphatase was optionally added, as indicated. The experiment shown is representative of two independently performed biological replicates.

## 3. Results

### 3.1 The TOPBP1-NBS1 interaction can be recapitulated using E. coli- and IVTT-produced proteins

Previous work from many different laboratories has shown that TOPBP1 and MRN act together to promote DDR signaling, and that NBS1 and TOPBP1 can bind to one another, although how this interaction occurs is not known (Morishima et al, 2007; Yoo et al, 2009; Duursma et al., 2013; Shiotani et al, 2013; Montales et al, 2022). The domain structures of the two proteins are shown in Figure 1A. As detailed above, TOPBP1 contains nine BRCT domains, named 0-8. NBS1 is less complex, with three protein-protein interaction motifs, two BRCT domains and one FHA domain. One previous study analyzed the interaction between TOPBP1 and NBS1 in some detail (Yoo et al, 2009). Using *Xenous* egg extracts, Yoo and colleagues found that TOPBP1 and NBS1 could co-precipitate under resting conditions, and that the interaction was enhanced when DDR signaling was activated. This work went on to show that TOPBP1 uses its BRCT0-2 region to bind NBS1, and that the relevant region within NBS1 mapped to the amino-terminal 400 amino acids of the protein. We have recently shown that MRN is important for recruitment of TOPBP1 in frog egg extracts to long, linearized dsDNA templates that mimic DSBs (Montales et al, 2022). Given these findings, we were motivated to analyze the TOPBP1-NBS1 interaction in more detail.

We first sought to repeat some of the earlier findings by Yoo and colleagues (Yoo et al, 2009), that a fragment of *Xenopus* NBS1 corresponding to amino acids 1-400 could bind to *Xenopus* TOPBP1 after incubation in frog egg extracts. Accordingly, we expressed and purified from *E. coli* a GST-NBS1 1-400 fusion protein. This protein, along with GST alone, was added to egg extract in either the presence or absence of linearized 5kb dsDNA, which we have previously shown will potently activate both ATR and ATM in our extracts (Montales et al, 2021; Ruis et al, 2022; Montales et al, 2022). The GST protein complexes were then recovered back out of the extract, using glutathione-sepharose beads, and the presence of endogenous TOPBP1 was detected via Western blotting. As displayed in Figure 1B, TOPBP1 did not bind to GST alone, but it did bind to GST-NBS1 1-400. Binding was evident in the sample lacking DSBs and was modestly enhanced by the presence of DSBs. We next asked if there are additional binding sites for TOPBP1 within NBS1, beyond those present in the 1-400 region, and we found that this is not likely as TOPBP1 did not bind to a GST fusion containing NBS1 from residue 393 to the end of the protein, amino acid 763 (Figure 1C). Lastly, we wanted to confirm that the BRCT0-2 region of TOPBP1 could bind to NBS1 1-400, and for this a myc-tagged TOPBP1 fragment corresponding to BRCT0-2 was produced by in vitro transcription and translation (IVTT) and added to egg extract along with GST-NBS1 1-400. DSBs were optionally added, and we observed that myc-TOPBP1 BRCT0-2 could readily bind to GST-NBS1 1-400, but not to GST alone, and that binding was modestly enhanced by DSBs (Figure 1D). Taken together, the data in Figures 1A-D are all in excellent agreement with the previous work (Yoo et al, 2009).

As shown here, and by others, a constitutive interaction between TOPBP1 and NBS1 occurs in egg extract even in the absence of DDR signaling (Figures 1B-D; Yoo et al, 2009). We wanted to study this basal interaction further, and we first asked if we could recapitulate it using a protein-binding assay that we nicknamed Bugs-Bunny (Figure 2A). In the Bugs-Bunny system one binding partner is expressed and purified from *E. coli* (Bugs) and the other is synthesized by IVTT in rabbit reticulocyte lysates (Bunny). After synthesis of one binding partner by IVTT we simply add the purified form of the other partner to the rabbit reticulocyte lysates. After incubation the samples are diluted in binding buffer and protein complexes isolated using an affinity resin corresponding to the tag on the *E. coli* produced protein (for this study the tag is predominantly GST, but sometimes MBP). This system has several advantages for the study of protein-protein interactions. One, the endogenous proteins present in the reticulocyte lysates help to prevent non-specific binding of the target protein to the resin. Two, as detailed below, the reticulocyte lysates can contain modifying enzymes, such as casein kinase 2 (CK2), that are important for certain interactions. Three, and most importantly, because synthesis of a target protein by IVTT is far less labor intensive than purification of the target from bacterial or insect cells, one can rapidly screen through a collection of mutants to analyze sequence determinants required for the interaction. Lastly, in our experience, proteins are much better behaved, in terms of proper folding and solubility, when made by IVTT relative to other expression systems.

We first examined binding of GST-NBS1 1-400 to myc-tagged TOPBP1 BRCT0-2 in the Bugs-Bunny assay. These are both derivatives of the *Xenopus* proteins, as are all the factors analyzed in this study. As shown in Figure 2B, myc-BRCT0-2 could bind to GST-NBS1 1-400 under these assay conditions, but not to GST alone. These data show that the NBS1::TOPBP1 interaction is readily observed using the Bugs-Bunny system, and thus that egg extract is not required for the interaction to occur. We next assessed the specificity of the interaction between NBS1 and TOPBP1’s BRCT0-2 region by asking if NBS1 could also bind to TOPBP1’s BRCT4&5 domains. Binding assays were run with the following partners: GST-NBS1 1-400 with myc-tagged TOPBP1 BRCT0-2 or 4&5 domains, and GST fused to the RAD9 Tail domain (RAD9Tail) with the same pair of TOPBP1 proteins. We included RAD9Tail because we had previously shown that it binds well to myc-BRCT0-2 in our Bugs-Bunny system (Kim et al, 2020). As shown in Figure 2C, myc-BRCT0-2 could bind both NBS1 and the RAD9Tail, and myc-BRCT4&5 failed to bind efficiently to either NBS1 or RAD9Tail. Quantification revealed that binding of myc-TOPBP1 BRCT0-2 to NBS1 was slightly more efficient than binding to RAD9Tail, and that binding of myc-BRCT4&5 to either GST substrate was largely undetectable. We conclude that the Bugs-Bunny system faithfully recapitulates a specific interaction between NBS1 and TOPBP1.

Both BRCT1 and BRCT2 from TOPBP1, as well as BRCT1 from NBS1, contain PBPs, suggesting that the TOPBP1::NBS1 interaction may be phospho-dependent. Previous work from our group has shown that the interaction between TOPBP1 and the RAD9Tail, which requires phosphorylation of RAD9Tail, can be recapitulated in the Bugs-Bunny system and this is because the reticulocyte lysates used for IVTT contain CK2, the kinase responsible for RAD9Tail phosphorylation (Kim et al, 2020). Thus, we next asked if including phosphatase in the binding assay would prevent the NBS1::TOPBP1 interaction, as it does the RAD9Tail::TOPBP1 interaction. Myc-BRCT0-2 was produced by IVTT and, prior to addition of the GST fusion proteins, the samples were optionally treated with lambda phosphatase. As expected, phosphatase treatment eliminated the RAD9Tail::TOPBP1 interaction, however the NBS1::TOPBP1 interaction was refractory to phosphatase treatment (Figure 2D). We conclude that the NBS1::TOPBP1 interaction does not require the presence of phosphate on either binding partner.

### 3.2 NBS1 can bind to both BRCT1 and BRCT2 of TOPBP1

To pursue these observations, we next asked if NBS1 required an intact BRCT0-2 region to associate with TOPBP1. To explore this, we subdivided TOPBP1’s BRCT0-2 region into the individual BRCT domains and tested them for binding to NBS1. Surprisingly, both BRCT1 and BRCT2 could bind to NBS1 (Figure 3A), whereas BRCT0 could not. In addition, while NBS1 readily binds the isolated BRCT1 or BRCT2 domains, the same is not true of RAD9Tail, as it could only interact with the contiguous BRCT0-2 region, and not the individual BRCT1 or BRCT2 domains (Figure 3B). Taken together, the data presented in Figures 1-3 show that the TOPBP1 BRCT0-2 region interacts with NBS1 in a fundamentally different manner than it does with RAD9Tail. The RAD9Tail::TOPBP1 interaction requires an intact BRCT0-2 region and is dependent on phosphorylation (summarized in Figure 3C), whereas the NBS1::TOPBP1 interaction requires neither an intact BRCT0-2 region nor phosphorylation (Figure 3D).

**Figure 3.**
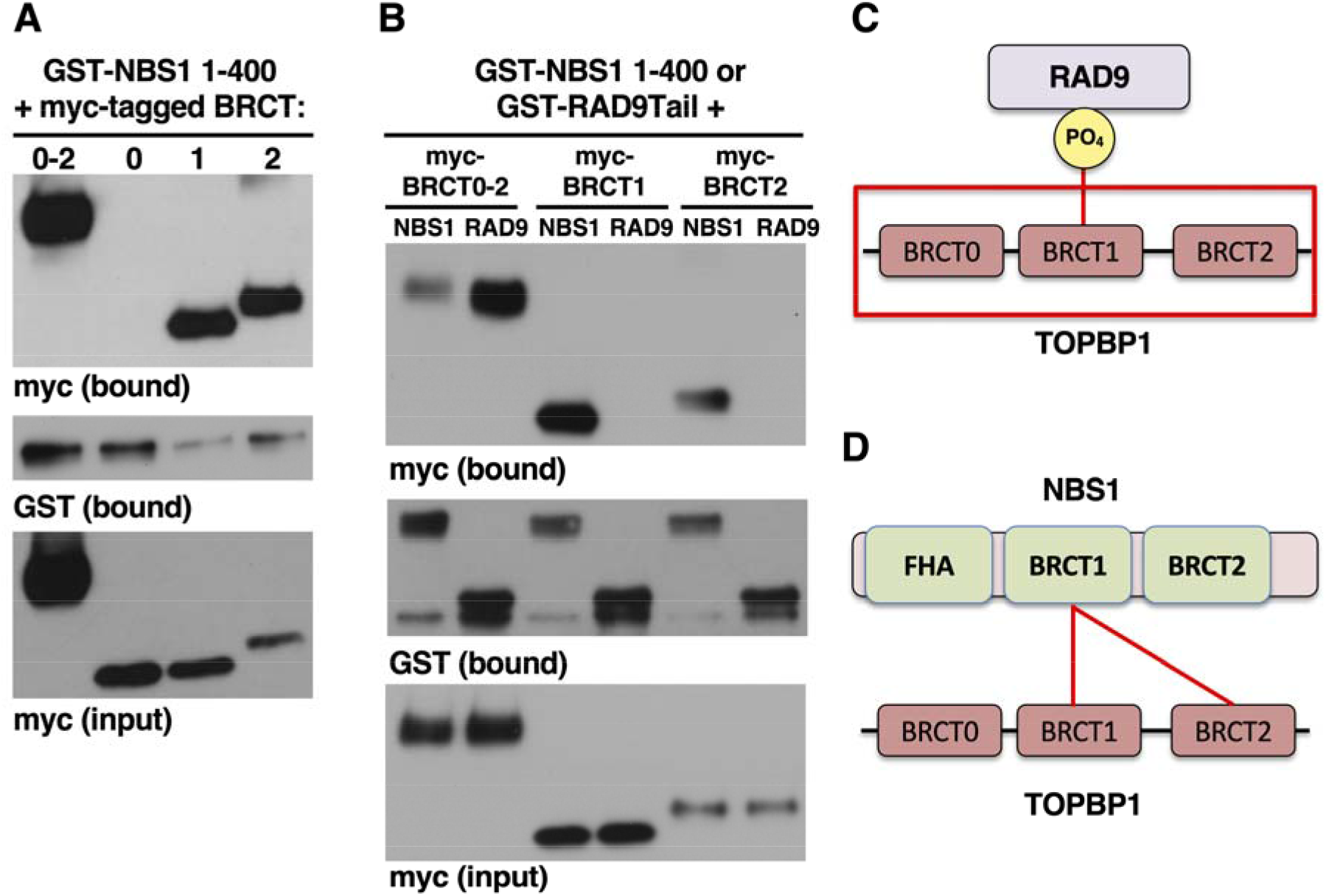
NBS1 independently binds to both BRCT1 and BRCT2 from TOPBP1. **A**. GST pull-down assays, using GST-NBS1 1-400 with myc-tagged versions of TOPBP1’s N-terminal BRCT domains, were performed exactly as in Figure 2B. **B**. GST pull-down assays, using GST-NBS1 1-400 or GST-RAD9Tail together with myc-tagged versions of TOPBP1’s N-terminal BRCT domains, were performed exactly as in Figure 2B. The experiments shown are representative of two independently performed biological replicates. **C-D**. Cartoons highlighting the different ways that RAD9Tail and NBS1 interact with TOPBP1’s BRCT0-2 region. See text for details.

### 3.3 NBS1 uses its BRCT1 domain to bind TOPBP1

We next examined the sequence determinants within NBS1 that allow binding to TOPBP1’s BRCT1&2 region. The constructs used for this are summarized in Figure 4A. We first confirmed that full-length NBS1 could bind TOPBP1’s BRCT1&2 domains in the Bugs-Bunny system, and to do so we employed an MBP-TOPBP1 BRCT1&2 fusion protein purified from *E. coli*, together with myc-tagged NBS1, made by IVTT. As shown in Figure 4B, full-length myc-NBS1 can bind MBP-TOPBP1 BRCT1&2, but not MBP alone. We also observed that MBP-TOPBP1 BRCT1&2 could bind the NBS1 1-400 region, but not the 393-763 region (Figure 4C). These data show that despite switching production of the TOPBP1 region from IVTT to *E. coli*, and vise-versa for NBS1, the ability to interact was not altered. We next used MBP-TOPBP1 BRCT1&2 to delineate the region within NBS1 to which it binds. The N-terminal portion of NBS1 was subdivided into three fragments, 1-100 (FHA domain), 100-200 (BRCT1), and 200-400 (BRCT2 plus additional sequences, see Figure 4A), and we observed that only the BRCT1 domain could bind to TOPBP1 BRCT1&2 (Figure 4D). Thus, unlike TOPBP1 where both BRCT1 and BRCT2 bind NBS1, for NBS1 just BRCT1 and not BRCT2 binds to TOPBP1.

**Figure 4.**
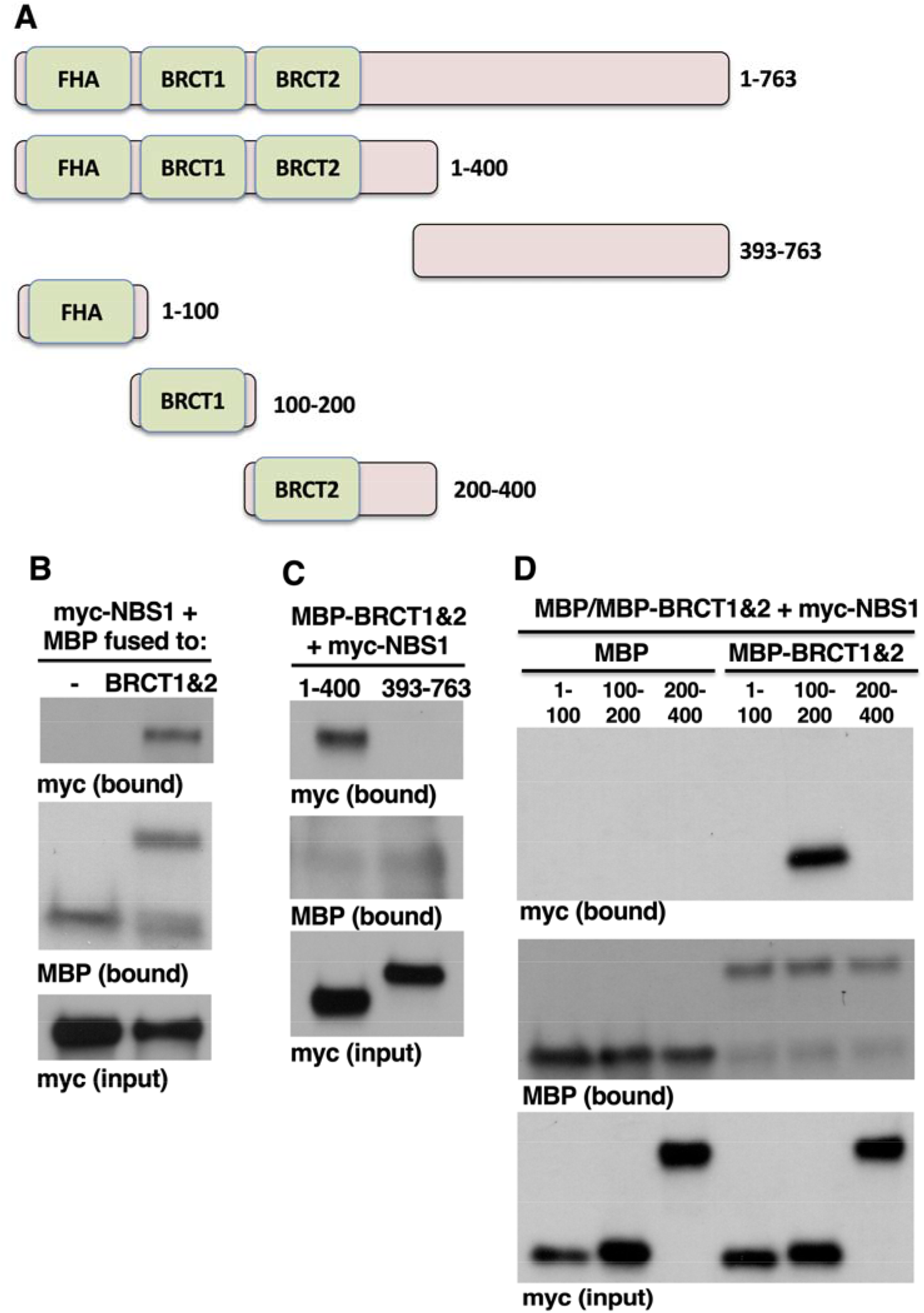
NBS1 uses its BRCT1 to bind TOPBP1. **A**. Cartoons summarizing the domains located within the N-terminus of NBS1 and the various truncations used in the binding assays. **B**. MBP pull-down assays were performed using either MBP alone or MBP fused to TOPBP1’s BRCT1&2 domains, together with myc-tagged and full-length NBS1. The experiment shown is representative of two independently performed biological replicates. **C**. MBP pull-down assays were performed using MBP-TOPBP1 BRCT1&2, together with the indicated myc-tagged forms of NBS1. The experiment shown is representative of two independently performed biological replicates. **D**. MBP pull-down assays were performed using either MBP alone or MBP fused to TOPBP1’s BRCT1&2 domains, together with the indicated myc-tagged forms of NBS1. The experiment shown is representative of two independently performed biological replicates.

### 3.4 Direct interaction between NBS1 and TOPBP1

An important question became is the NBS1::TOPBP1 interaction direct, or does it require a bridging factor that might be present in the reticulocyte lysates used for our binding assays? To answer this we expressed and purified an MBP-NBS1 1-200 fusion protein from *E. coli* (Figure 5A). We attempted to purify MBP-NBS1 100-200, however we found that this protein was insoluble and difficult to express in *E. coli*. We also produced a 6His-T7-TOPBP1 BRCT0-2 recombinant protein (Figure 5A), one that we have used in the past (Kim et al, 2020). Thus, in this experiment, both proteins were expressed in *E. coli* and then purified, thereby eliminating concerns about bridging factors. To block non-specific binding to the resin the assays were performed in binding buffer supplemented with 1% BSA. As shown in Figure 5B, 6His-T7-TOBP1 BRCT0-2 could readily bind MBP-NBS1 1-200, but not MBP alone, and binding was not inhibited by inclusion of phosphatase. We conclude that the interaction between NBS1 and TOPBP1 is direct and does not require any post-translational modifications on either partner.

**Figure 5.**
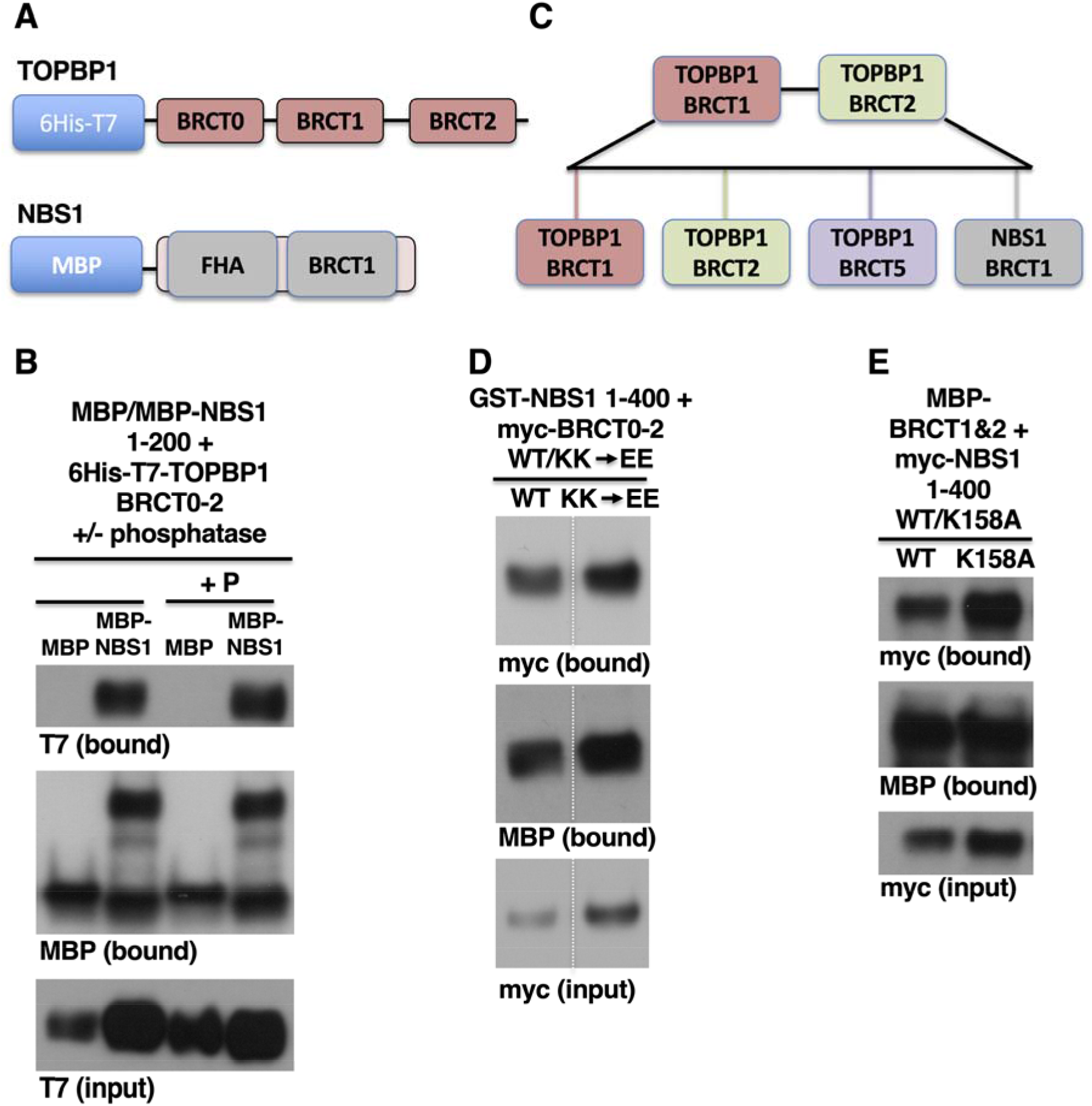
Direct interaction between TOPBP1 and NBS1 and the role of PBPs in the interaction. **A**. Cartoons showing the recombinant proteins used in the binding assay displayed in part B. **B**. All the proteins used in this binding assay were expressed and purified from *E. coli. Shown is* an MBP pull-down assay with either MBP alone or MBP fused to NBS1 1-200, together with 6His-T7 tagged TOPBP1 BRCT0-2. The binding reaction was performed in the presence of a 1% BSA solution, to suppress non-specific binding to the maltose resin. As in Figure 2D, phosphatase was optionally included (as indicated by the “P”). The experiment shown is representative of two independently performed biological replicates. **C**. A schematic summarizing the disparate BRCT domains that are known to bind the TOPBP1 BRCT1&2 region. All of these domains contain a PBP. D. GST pull-down assays, using GST-NBS1 1-400 with myc-tagged TOPBP1 BRCT0-2, were performed exactly as in Figure 2B. Shown are both the wild type (WT) and K155K249 to EE (“KK->EE”) forms of TOPBP1 BRCT0-2. The experiment shown is representative of two independently performed biological replicates. We note that the image shown is derived from a larger gel, and the irrelevant samples were removed. The white dashed line depicts where the gel was spliced together. The full gel with all original samples is shown in Figure S1B. **E**. An MBP pull-down assay was performed with MBP-TOPBP1 BRCT1&2 together with either wild type (“WT”) or the lysine 158 to alanine (“K158A”) forms of myc-tagged NBS1. 1-400. The experiment shown is representative of two independently performed biological replicates.

### 3.5 No requirement for phosphate-binding pockets in the TOPBP1::NBS1 interaction

Our recent study (Kim et al, 2020), together with data shown here, have demonstrated that TOPBP1’s BRCT1&2 region interacts with several different BRCT domains, including TOPBP1’s BRCT1, BRCT2, and BRCT5 domains as well as BRCT1 from NBS1 (Figure 5C). One feature that all these domains have in common is a PBP. Even though we have shown that phosphate *per se* is not needed for the TOPBP1::NBS1 interaction (Figures 2D and 5B), it was still possible that the PBPs played a role in the binding. To test this idea we produced mutant proteins where a key and highly conserved lysine residue within the PBP was mutated. For TOPBP1 we mutated both K155 in BRCT1 and K249 in BRCT2 to glutamic acid, in the context of a myc-tagged BRCT0-2 fragment. Previous work with human TOPBP1 has shown that a similar K to E substitution in BRCT1 prevents binding to RAD9 (Rappas et al, 2011), however we observed that the KK155,249EE double mutant could still readily bind to GST-NBS1 1-400 (Figure 5D). For NBS1, we initially made the same K to E mutation at K158 in BRCT1 and observed that this mutant could not bind to TOPBP1 (data not shown). Because this charge-reversal mutation might non-specifically disrupt folding of the NBS1 domains we also tested a more subtle mutation, substituting K158 with alanine. The K158A mutation is still expected to inactivate the PBP, as has been shown for the BRCA1 PBP (Botuyan et al, 2004), however we observed that it had no impact on binding to TOPBP1 (Figure 5E). Thus, for both TOPBP1 and NBS1, an intact PBP is dispensable for their interaction.

### 3.6 Intact BRCT domains are not required for the TOPBP1::NBS1 interaction

To learn more about the NBS1::TOPBP1 interaction we asked how creating truncations of TOPBP1’s BRCT0-2 region would impact binding to NBS1 1-400. A series of deletion mutants were constructed as fusions to MBP and are shown in Figure S2A. These MBP fusions were produced by IVTT and tested for binding against GST-tagged NBS1 1-400. We found that MBP-BRCT0-2 (amino acids 1-333) and MBP-BRCT1&2 (amino acids 91-275) could bind NBS1 (Figure S2B). We next asked if the linker sequence between BRCTs 1 and 2 influenced binding of BRCT1, by comparing a construct containing the linker (amino acids 91-196) to one that does not (amino acids 91-179). As shown in Figure S2B, both proteins could bind NBS1. Moving forward, we next asked if BRCT1 must be in an intact form to bind to NBS1. Structural analysis of BRCT domains has shown that they are highly ordered, using a series of alpha helices and beta sheets to fold into a unit with a conserved topology (reviewed by Gerloff et al, 2012; Leung and Glover, 2013; Reinhardt and Yaffe, 2013; Wan et al, 2016). Thus, our expectation was that by making deletions within BRCT1 the domain would lose its ability to fold properly and binding to NBS1 would be lost. This, however, was not the case. The Conserved Domain sequence analysis tool that is available on the NCBI Structure website (www.ncbi.nlm.nih.gov/Structure/cdd/wrpsb.cgi) defines *Xenopus* TOPBP1 BRCT1 as running from amino acid 108 to 179. As shown in Figure 6A&B, and we observed that a fragment containing amino acids 128-179, and therefore missing the N-terminal 20 amino acids of BRCT1, could bind well to NBS1. A slightly smaller piece, corresponding to amino acids 138-179, could not bind NBS1 (Figure 6B). Intrigued by these observations we next asked if loss of C-terminal sequences within TOPBP1’s BRCT1 would impact binding to NBS1. We found that truncation of the C-terminal 23 amino acids of BRCT1, using a construct containing amino acids 91-156, did not disrupt binding and, furthermore, that an even smaller fragment, amino acids 128-156, could bind efficiently to NBS 1 1-400 (Figure 6C). Thus, an intact BRCT1 domain is not required for its interaction with NBS1. To see if this is also true of TOPBP1’s BRCT2 we generated a series of deletion constructs (Figure S2C) and tested them for binding to GST-NBS1 1-400. These results showed that a small fragment of 23 amino acids, corresponding to residues 250-273, was sufficient for binding to NBS1 (Figures S2D&E). We conclude that small sequences within both BRCT1 and BRCT2 of TOPBP1 can confer binding to NBS1 onto a heterologous protein, in this case MBP.

**Figure 6.**
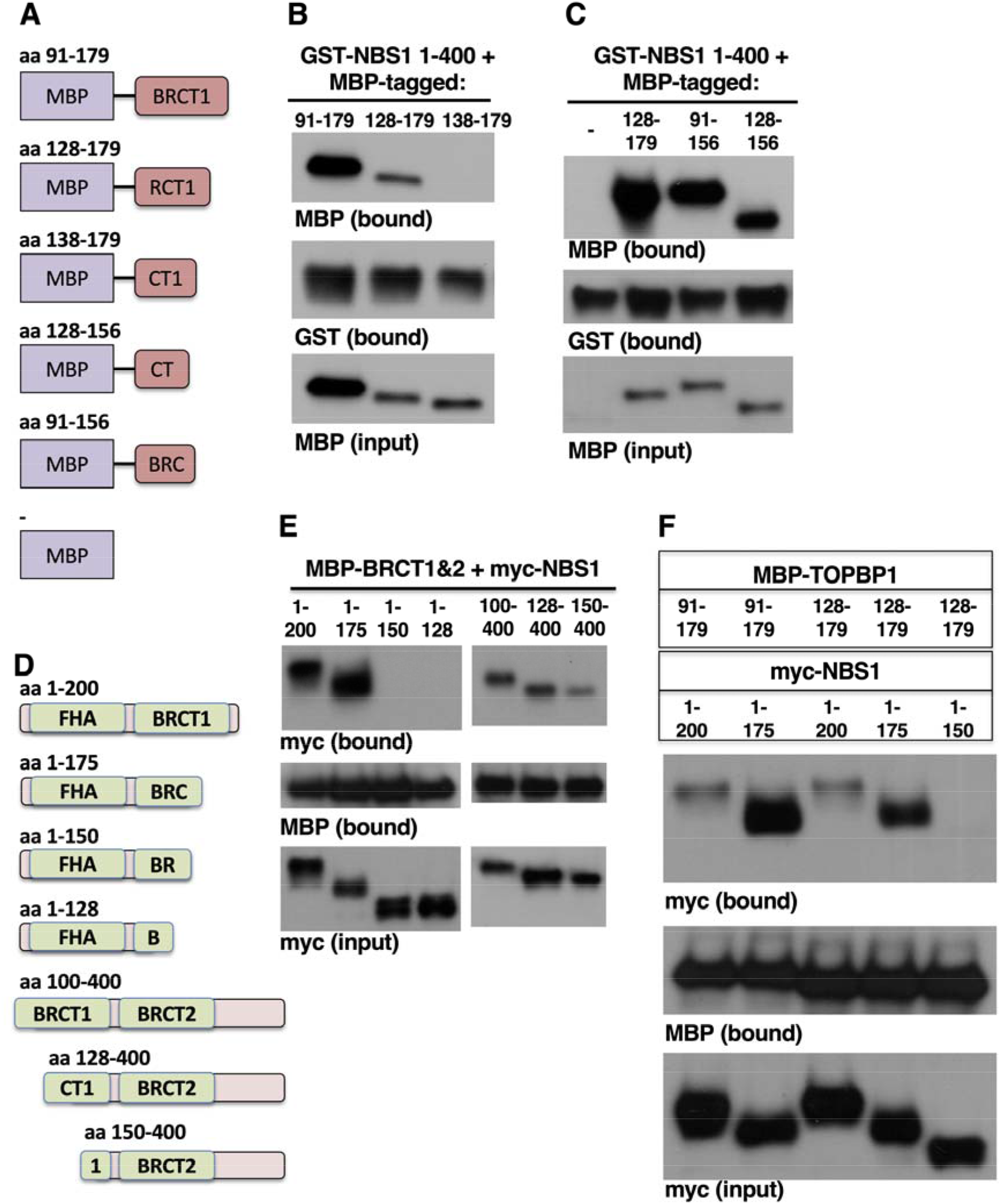
Intact BRCT domains are dispensable for the TOPBP1::NBS1 interaction. **A**. Cartoons summarizing the various TOPBP1 BRCT1 truncations used in the binding assays. These MBP-fusion proteins were produced by IVTT and used in pull-down assays with recombinant GST-NBS1 1-400. **B-C**. GST pull-down assays, using GST-NBS1 1-400 together with the indicated MBP-tagged TOPBP1 derivatives. The experiments shown are representative of two independently performed biological replicates. **D**. Cartoons summarizing the various NBS1 truncations used in the binding assays. E. MBP pull-down assays using MBP-TOPBP1 BRCT1&2 together with the indicated myc-tagged NBS1 derivatives. The experiment shown is representative of two independently performed biological replicates. **F**. MBP pull-down assays were performed with the indicated proteins. In this experiment, both the MBP-TOPBP1 derivatives and the myc-NBS1 derivatives were produced by IVTT. The experiment shown is representative of two independently performed biological replicates.

We next examined sequence determinants within the NBS1 BRCT domain that allow binding to TOPBP1. For this we generated the series of deletion mutants shown in Figure 6D, and we tested them for binding to *E. coli*-produced MBP-TOPBP1 BRCT1&2. As expected, we found that the fragment corresponding to intact FHA and BRCT1 domains (1-200) could bind efficiently to TOPBP1 (Figure 6E). We also observed that a fragment where the carboxyl terminal 25 amino acids of BRCT1 were removed (1-175) retained the ability to bind TOPBP1. Thus, like TOPBP1’s BRCT1&2 domains, NBS1’s BRCT1 domain can be truncated without attenuating the interaction. We then asked if removing more of the carboxyl terminal sequences impacted binding and found that neither a fragment corresponding to amino acids 1-150 nor one corresponding to 1-128 could bind TOPBP1. We also examined amino-terminal deletions and found that removing the amino-terminal half of the NBS1 BRCT1 domain did little to inhibit the interaction with TOPBP1 (Figure 6E). These data show that, like TOPBP1’s BRCT1 and BRCT2 domains, the NBS1 BRCT1 domain need not be in an intact and properly folded configuration to bind TOPBP1.

Data presented thus far show that, for all the relevant BRCT domains in this study, binding persists even after truncation of the domain. These curious results inspired us to ask if the interaction required that at least one partner contain an intact BRCT domain. For example, if the NBS1 BRCT1 domain and the TOPBP1 BRCT1 domain are both in a truncated form, would the interaction be lost? As it turned out, the answer is no. For this experiment we used an assay system where both binding partners were produced by IVTT. This was because all the required constructs for this experiment were available for protein production by IVTT, and thus there was no need to remake them for expression in *E. coli*. Proteins were produced by IVTT and then the lysates were mixed, to allow interaction. The samples were then diluted in binding buffer and incubated with maltose beads, followed by washing, elution, and Western blotting. As shown in Figure 6F, binding was readily observed when the BRCT1 domains for both TOPBP1 and NBS1 were intact (lane 1). Similarly, binding was observed when TOPBP1 BRCT1 was intact and NBS1 BRCT1 was truncated (lane 2), and vise versa (lane 3). These results are all in accordance with the data derived from the Bugs-Bunny assays shown in Figures 6B, C, and E. When both binding partners where in a truncated form we also observed efficient binding (lane 4), and binding was not observed when the NBS1 derivative was truncated further (lane 5), consistent with data shown in Figure 6E. Based on these data, we conclude that neither the TOPBP1 BRCT1 domain nor its counterpart in NBS1 need to be in an intact and properly folded configuration for the interaction to occur.

## 4. Discussion

In this study we examined how two crucial DDR proteins, TOPBP1 and NBS1, interact with one another. Prior to this work, it was known that TOPBP1 and NBS1 can form a complex, in a manner conserved from frogs to man (Morishima et al, 2007; Yoo et al, 2009). However, it was not known which domains within either protein control the interaction, and neither was it known if the interaction was direct or mediated by a bridging factor. Here, we show that the NBS1 BRCT1 domain binds both the BRCT1 and BRCT2 domain from TOPBP1. These are specific interactions, as revealed by our finding that NBS1 binds well to TOPBP1 BRCTs 1&2 but shows very little affinity for BRCTs 4&5 (Figure 2C). Likewise, TOPBP1 binds NBS1 BRCT1 efficiently but there is no detectable binding to either the FHA or BRCT2 domains from NBS1 (Figure 4D). Our data also show that the interaction between TOPBP1 and NBS1 is constitutive, and it is also direct and independent of any post translational modifications (Figure 5). Thus, at least a subset of MRN complexes contain TOPBP1 even under resting conditions. In our previous work we showed that immunodepletion of MRN from frog egg extracts with antibodies targeting NBS1 co-depletes all detectable MRE11 but does not influence the levels of TOPBP1 (Montales et al, 2022). This shows that the association of TOPBP1 with NBS1 in egg extract is not as strong as the association of NBS1 with other members of MRN, although it is possible that the antibodies used for depletion dislodge TOPBP1 from NBS1.

What might be the function of the TOPBP1::NBS1 interaction(s)? In addition, a related question is why did TOPBP1 evolve two disparate binding sites for NBS1, positioned closely together at the amino terminus of the protein? We propose that one function of the interaction is to recruit TOPBP1 to sites of damage so that it can activate the ATR kinase. Three previous studies, all performed with frog egg extracts, have implicated MRN in recruitment of TOPBP1 to stalled replication forks (Lee and Dunphy, 2013), to an ATR-activating DNA structure comprised of M13 ssDNA to which primers are annealed (Duursma et al, 2013), and to 5kb linear dsDNAs that mimic DSBs (Montales et al, 2022). Thus, it is clear that MRN is important for how TOPBP1 arrives at sites of damage, and we propose that at least one of the interactions defined in this paper plays a role in TOPBP1 recruitment for the purpose of ATR signaling. But why does have TOPBP1 harbor more than one site for interaction with NBS1? One possibility is that TOPBP1 also collaborates with MRN is the physical repair of a DSB. Indeed, a recent study using Repair-seq to order DNA repair proteins into common pathways showed that TOPBP1 clusters with NBS1, MRE11, and RAD50 in that all four show a common repair defect upon their depletion (Hussmann et al, 2021). Thus, it may be that TOPBP1 interacts with NBS1 differently for ATR signaling versus DNA repair, and if so, this would explain the multiple binding sites. Clearly, further work, combining mutagenesis with functional assays, is needed to clarify the roles of the two NBS1 binding sites on TOPBP1.

Another important question raised by our studies is how are the BRCT domains from NBS1 and TOPBP1 interacting with one another? There is precedent for BRCT domains to form both homo- and heterodimers. The best studied example of this is found with the XRCC1 and LIG3 proteins (reviewed most recently by London, 2020). XRCC1 has two BRCT domains, with BRCT2 being the one that binds to itself as well as to the single BRCT domain within LIG3, which also interacts with itself. Our previous work has shown that TOPBP1’s BRCT1 and 2 domains can interact with one another, and we have shown here that they also bind to NBS1 BRCT1. Thus, the TOPBP1::NBS1 system is similar to XRCC1::LIG3 in that both systems can form homo- and heterotypic interactions. Recent structural analysis of a human XRCC1 BRCT1::LIG3 BRCT dimer revealed that the two BRCT domains bind one another largely through electrostatic interactions between residues located in the alpha helix 1 portion of the respective domains (Hammel et al, 2021). Thus, it will be of interest in future work to determine if a similar molecular interface controls the interactions between TOPBP1 and NBS1.

Work presented here also reveals the remarkable versatility that TOPBP1’s BRCT0-2 region displays in its interactions with various binding partners. On the one hand, there is the well-studied, phosphate-dependent interaction with RAD9Tail (and other phospho-substrates), and on the other hand, as we have shown here, is a very different mode of interaction, involving phosphate-free and direct binding to NBS1. There are important differences between the phosphate-dependent and phosphate-free modes of binding. For the former, a contiguous BRCT0-2 module is required for efficient binding. In the case of the RAD9Tail, previous work has shown that it is BRCT1 that contacts phosphate (Rappas et al, 2011), however BRCT1 alone is not sufficient for the interaction (Figure 3B), and in data not shown we have observed that BRCT0 also plays a positive role, as the BRCT0-2 region binds RAD9Tail more efficiently than does the BRCT1&2 region. Thus, the BRCT0-2 continuum is needed for optimal binding in the phosphate -dependent mode, while the phosphate-free mode can occur with isolated domains.

Another unexpected finding reported in this work is that interaction between the TOPBP1 and NBS1 BRCT domains can occur in the context of truncated, and thus improperly folded, domains. As shown in Figures 6B&C, for TOPBP1’s BRCT1, we used MBP fusions to ask if truncation of the intact domain would impede binding and found that it did not. Indeed, we were able to show that a small peptide of just 29 amino acids can confer the ability to bind NBS1 when grafted on to MBP. Similarly, we found that an even smaller peptide from TOPBP1’s BRCT2 domain has the same properties (Figures S2D&E). For NBS1, we found that removal of large portions of BRCT1 did not impair binding to TOPBP1 (Figure 6E). In Figure 7A the amino acid sequences of the three BRCT domains studied here are shown. For TOPBP1, the residues that allow biding to NBS1, in a transferrable manner, are highlighted in red. Also highlighted in red are the residues within NBS1 that represent the minimal region sufficient for binding to TOPBP1, as defined by deletion analysis (Figure 6E). We do not yet know if this sequence is transferrable, but this seems likely. Also shown is the canonical arrangement of beta sheets and alpha helices that promote proper folding of BRCT domains. While the minimally defined region within BRCT2 shows good overlap with the corresponding sequence from NBS1, there is very poor overlap between the two TOPBP1 sequences, perhaps suggesting that TOPBP1’s BRCT1 uses a different binding mode to interact with NBS1 than does its BRCT2 domain.

**Figure 7.**
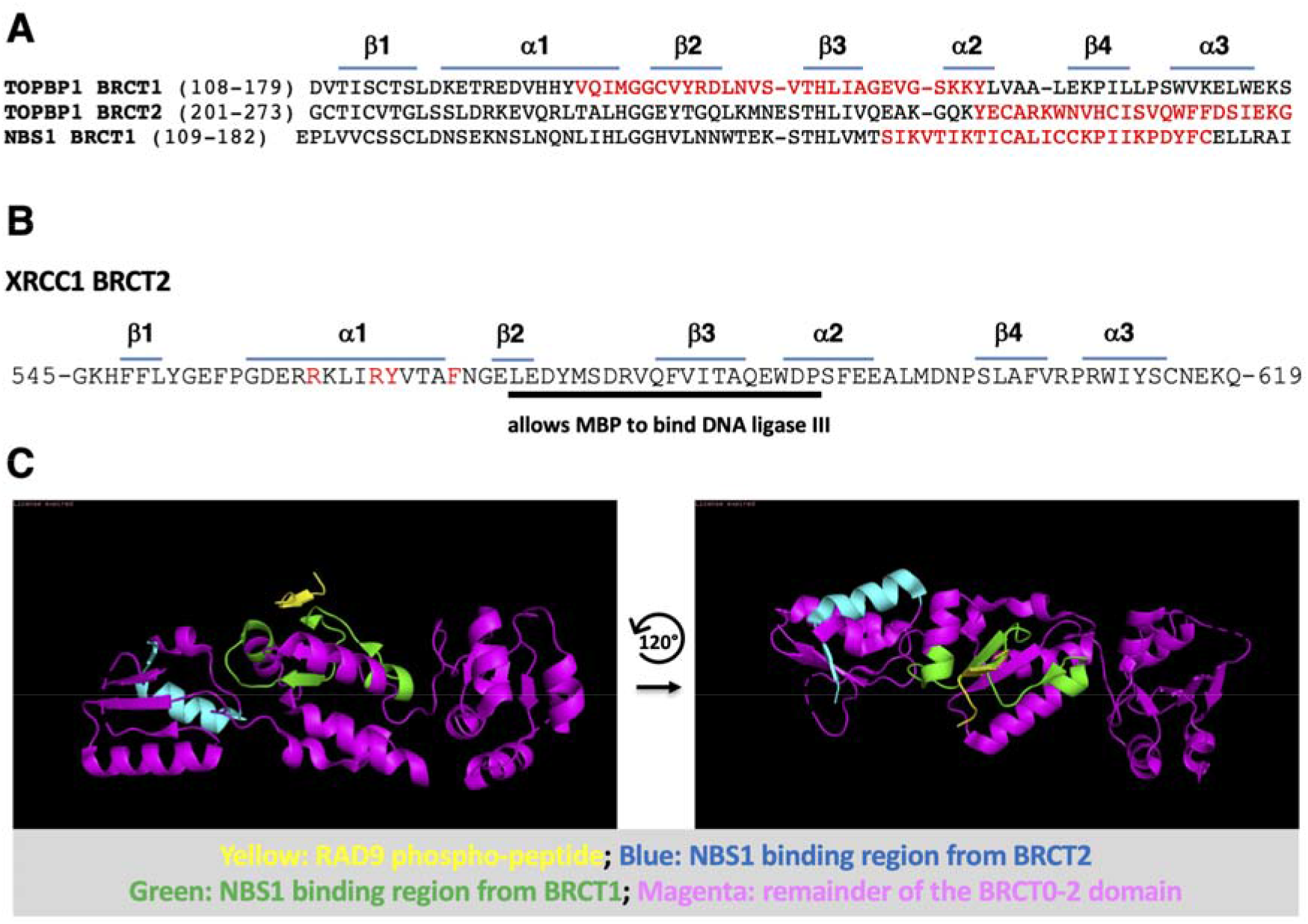
Summary of the BRCT domain delineation analyses. **A**. The amino acid sequences of the indicated domains are displayed. Above the sequences are the positions of either beta sheets or alpha helices, as indicated. Highlighted in red for TOPBP1 are the small stretches of amino acids that confer binding to NBS1 when fused to MBP. Highlighted in red for NBS1 is the region of BRCT1 that binds TOPBP1, as defined by deletion analysis. **B**. The human XRCC1 BRCT2 sequence is shown. Underlined in black is the small region that confers binding to LIG3 when fused to MBP, as shown in Taylor et al, 1998, and highlighted in red are residues shown by Hammel et al, 2021 to be required for binding to LIG3. **C**. Two views of the crystal structure of the human TOPBP1 BRCT0-2 region in complex with a RAD9 phospho-peptide. The images were generated using Pymol structure view software and data set PDB ID: 6HM5.

Given these results with truncations we went on to ask if binding could still occur when both partners where in a truncated form. We found that it could as truncated TOPBP1 BRCT1 clearly interacts with a truncated form of NBS1 BRCT1 (Figure 6F). This interaction is specific as the NBS1 1-175 truncation binds both intact TOPBP1 BRCT1 (91-179) and the truncated form (128-179), but further truncation of NBS1 to yield 1-150 yields a protein unable to bind either intact or truncated TOPBP1 BRCT1 (Figure 6F). Thus, truncated TOPBP1 BRCT1 binds to NBS1 with the same specificity as the intact form. While these results are difficult to reconcile with the notion that proper folding of an intact BRCT domain is a prerequisite for its ability to specifically interact with any given binding partner, they are not without precedent. Previous work has shown that a truncated form of XRCC1 retains the ability to bind to LIG3 (Taylor et al, 1998). Indeed, this work showed that a small peptide of 20 amino acids can be transferred to MBP, and this confers binding to LIG3. Furthermore, binding of the peptide to LIG3 was stoichiometric and of equivalent efficiency as the full-length XRCC1 protein (Taylor et al, 1998). The sequence of XRCC1 BRCT2 is shown in Figure 7B, and the LIG3-binding peptide underlined in black. Interestingly, this peptide maps to a similar position within its BRCT domain as does the peptide from TOPBP1’s BRCT1 that confers binding to NBS1, as both sequences are comprised of the residues that form beta sheet 3 within an intact domain, along with some flanking sequences (compare Figures 7A and 7B). Curiously, however, recent structural studies of the XRCC1::LIG3 complex have defined electrostatic interactions between residues largely located in alpha helix 1 as comprising the binding interface for the XRCC1::LIG3 interaction (Hammel et al, 2021). Thus, it is currently unclear how the XRCC1 20mer peptide maintains key properties of the interaction with LIG3 when the structural studies suggest the binding interface resides elsewhere in the domain. This question is relevant to the work presented here, given that small peptides from TOPBP1 can also confer binding to NBS1.

Lastly, we note that the crystal structure of human TOPBP1 in complex with a phospho-peptide from RAD9 has been solved (Day et al, 2018), and thus it was of interest to visualize where in this structure the two NBS1-binding peptides reside. As shown in Figure 7C, left panel, much of the NBS1 binding peptide from TOPBP1’s BRCT1 is solvent-exposed and overlaps with the region that binds RAD9 (in Figure 7C the NBS1-binding peptide is colored in green and the RAD9 peptide is colored in yellow). For the NBS1-binding peptide from BRCT2 we see that it is also solvent-exposed and thus, in principle, able to interact with heterologous proteins (Figure 7C, right panel). Having defined these two regions of TOPBP1 as able to interact with NBS1, the next important steps will be to first map the amino acids that are crucial for the interaction and to then test these mutants in functional assays for ATR activation and DNA repair, and such experiments are now in progress.

## Author contribution

W.M.M. conceptualized the project, acquired the funding, and wrote the manuscript together with O.H. O.H., K.R., and K.M. performed the experiments.

## Acknowledgments

This work was supported by NIH grant R01GM12287. We thank Karlene Cimprich for the gift of the GST-RAD9 Tail expression vector.

