## Supplementary material for "NBS1 binds directly to TOPBP1 via disparate interactions between the NBS1 BRCT1 domain and the TOPBP1 BRCT1 and BRCT2 domains": Supplemental Information.docx

**Supplemental Figures and Legends**

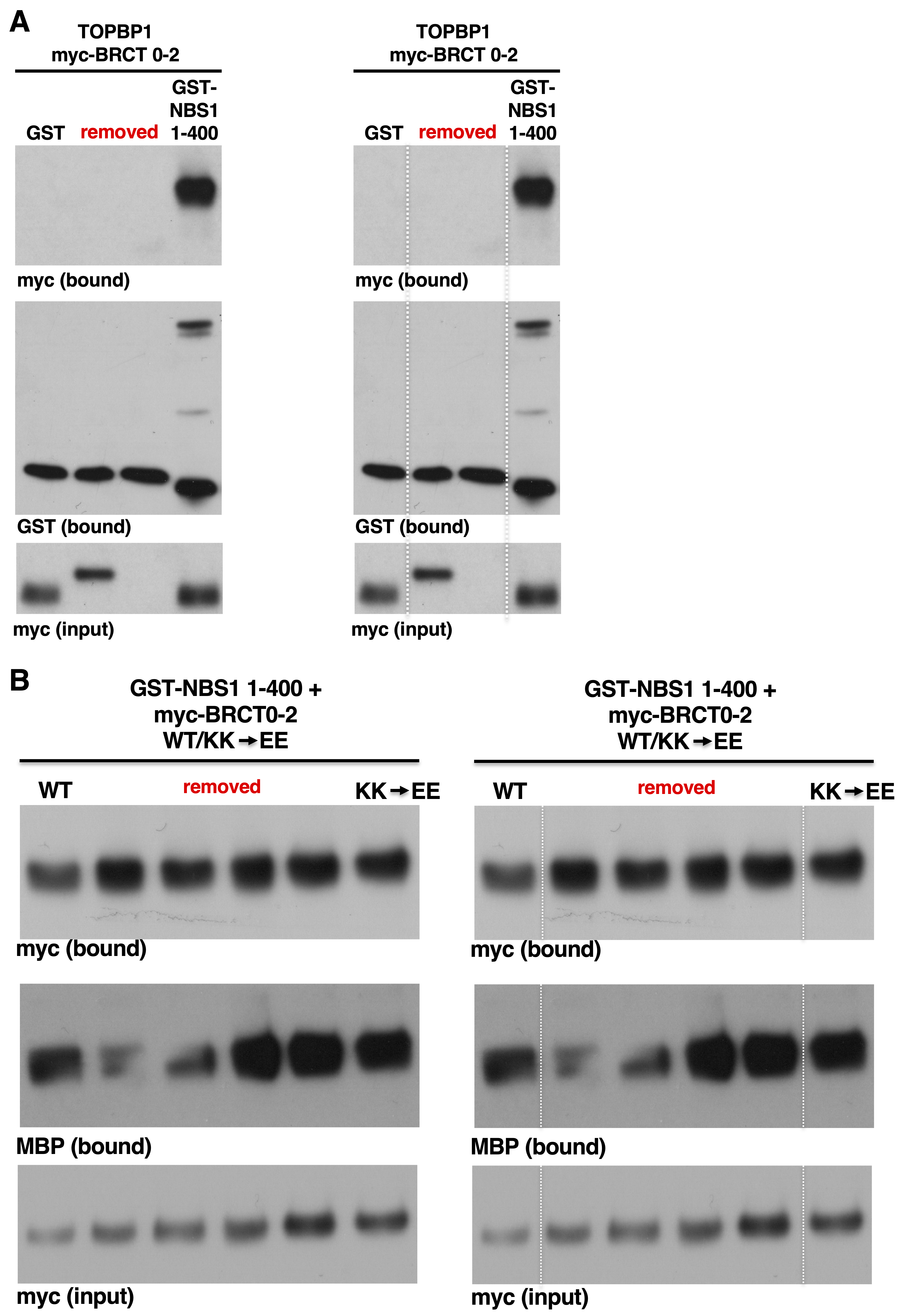

**Figure S1. Full blots for the experiments shown in the main figures 2B and 5D**

**A.** The full, unspliced blot for main figure 2B is shown, both with and without the dashed lines. **B.** The full, unspliced blot for main figure 5D is shown, both with and without the dashed lines.

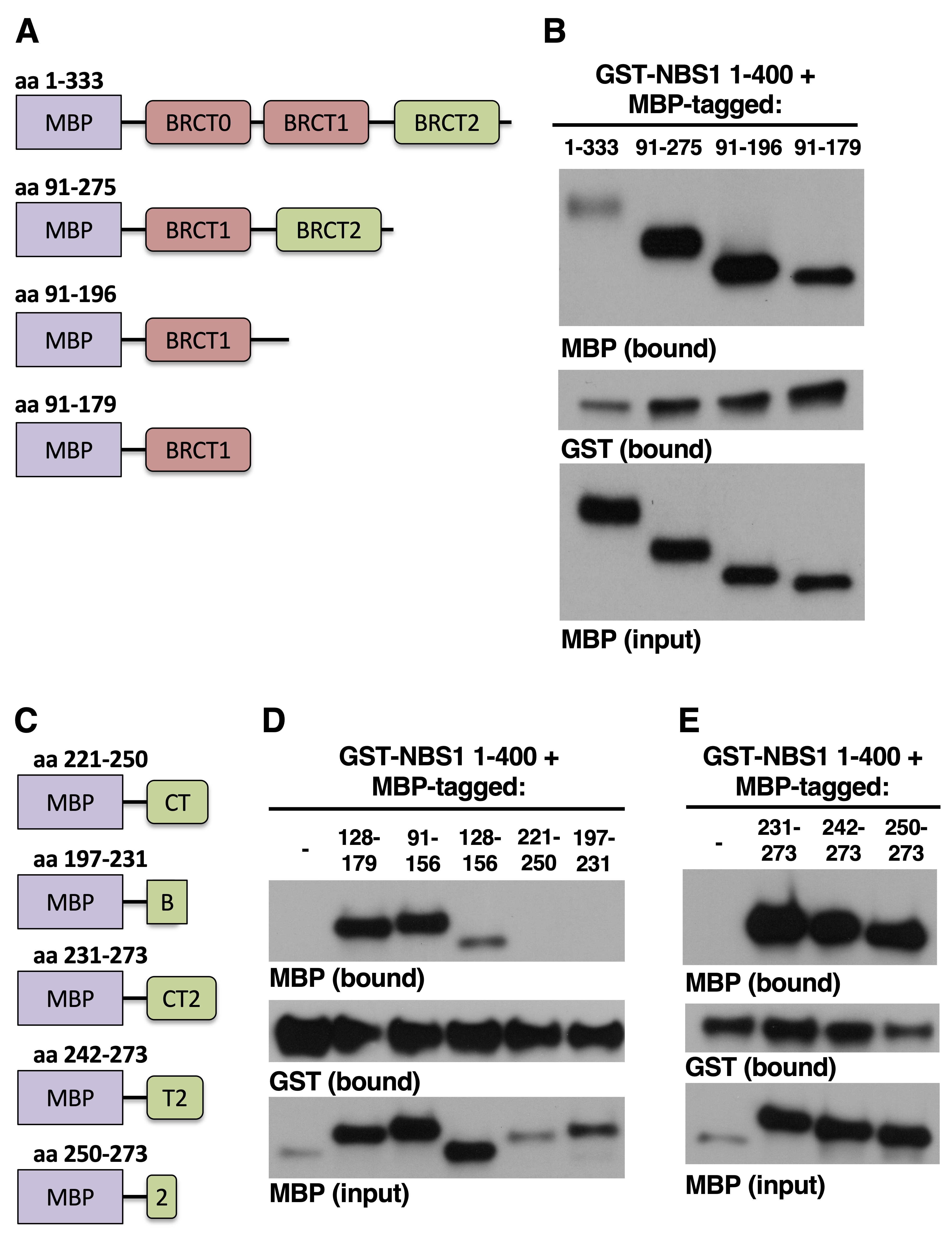

**Figure S2. Deletion analysis of TOPBP1 BRCT2 binding to NBS1.**

**A.** Cartoons showing the MBP fusions used in the deletion analysis for part B. **B.** A GST pull-down assay with GST-NBS1 1-400 and the indicated MBP-tagged fragments of TOPBP1. **C.** Cartoons showing the MBP fusions used in the deletion analysis for part D. **D.** A GST pull-down assay with GST-NBS11-400 and the indicated MBP-tagged fragments of TOPBP1 BRCT2.
